# ZYS-1 is not an ADAR1 inhibitor

**DOI:** 10.1101/2025.08.08.669362

**Authors:** Cassandra N. Smoak, Estelle N. Gardner, Renee N. Chua, Kyle A. Cottrell

## Abstract

Adenosine deaminase acting on RNA 1 (ADAR1) edits double-stranded RNA (dsRNA) substrates by the deamination of adenosine to inosine in a process known as A-to-I editing. Modulation of ADAR1 expression and editing activity has previously been described to play a role in cancer development and progression, with upregulation of ADAR1 being observed in a range of cancers. Further, depletion of ADAR1 leads to increased sensing of endogenous dsRNAs by dsRNA sensors in cell lines that require ADAR1 for survival, which are termed ADAR1-dependent. The activation of these sensors induces downstream production of type I interferons as well as translational inhibition and apoptosis. Therefore, ADAR1 is a promising oncologic therapeutic target. Recently, the small molecule ZYS-1 has been developed and presented as a direct inhibitor of ADAR1. We performed a series of in vitro and cellular experiments to validate the efficacy and specificity of ZYS-1 as an ADAR1 inhibitor. Evaluating the effect of ZYS-1 on cell viability revealed it to be equally cytotoxic to both ADAR1-dependent and ADAR1-independent cell lines, as well as wildtype and ADAR1 knockout cells. Moreover, ZYS-1 treatment had little effect on activation of PKR or induction of IFN stimulated genes. Importantly, treatment with ZYS-1 did not reduce cellular A-to-I editing for several known ADAR1 editing sites, and did not inhibit in vitro A-to-I editing by recombinant ADAR1. Together, these data indicate that ZYS-1 is not a selective inhibitor of ADAR1.

## Introduction

The RNA editing enzyme ADAR1 is an important regulator of double-stranded RNA (dsRNA) sensing pathways (Cottrell et al. 2024a; Heraud-Farlow and Walkley 2020). Adenosine-to-inosine (A-to-I) editing by ADAR1 prevents activation of MDA5 by endogenous dsRNAs (Liddicoat et al. 2015; Pestal et al. 2015; Mannion et al. 2014). Additionally, ADAR1 suppresses activation of the dsRNA sensor PKR in an RNA editing-independent manner (Sinigaglia et al. 2024; Chung et al. 2018; Hu et al. 2023). These functions of ADAR1 are essential in embryonic development and, importantly, in many cancer cell lines (Wang et al. 2004; Hartner et al. 2004; Gannon et al. 2018; Liu et al. 2019; Kung et al. 2021). Depletion of ADAR1 in roughly half of cancer cells causes cell death (Liu et al. 2019; Gannon et al. 2018; Kung et al. 2021). In these ADAR1-dependent cells, depletion of ADAR1 causes activation of PKR – which drives cell death – and activation of MDA5 leading to type I interferon signaling and expression of IFN-stimulated genes (ISGs). The IFN response caused by ADAR1 knockout can overcome resistance to immune checkpoint blockade (Ishizuka et al. 2019). These phenotypes associated with ADAR1 depletion make ADAR1 an important therapeutic target for many cancers, especially those with poor response to immune checkpoint blockade. However, there are currently no FDA-approved ADAR1 inhibitors.

The substrate of ADAR1 is an adenosine residue within duplexed RNA (Bass and Weintraub 1988; Wagner and Nishikura 1988). While the modified adenosine can either be base-paired or not (Wong et al. 2001), ADAR1 has no activity on single-stranded RNA (Bass and Weintraub 1988; Wagner and Nishikura 1988), or on adenosine or adenine alone (Polson et al. 1991). Previously, two small molecule adenosine analogues have been reported to disrupt ADAR1 function – 8-azaadenosine (Ramírez-Moya et al. 2020; Zipeto et al. 2016) and 8-chloroadenosine (Ding et al. 2020) – however, through extensive evaluation, we found that neither are selective inhibitors of ADAR1 (Cottrell et al. 2021). Recently, ZYS-1, another adenosine analogue (Figure 1a), has been reported to inhibit ADAR1(Wang et al. 2025) and has been used in subsequent studies (Shi et al. 2025). Here we carefully evaluate the efficacy and specificity of ZYS-1 as an ADAR1 inhibitor. Through a series of cellular and in vitro experiments, we found no evidence supporting ZYS-1 as an ADAR1 inhibitor.

**Figure 1.**
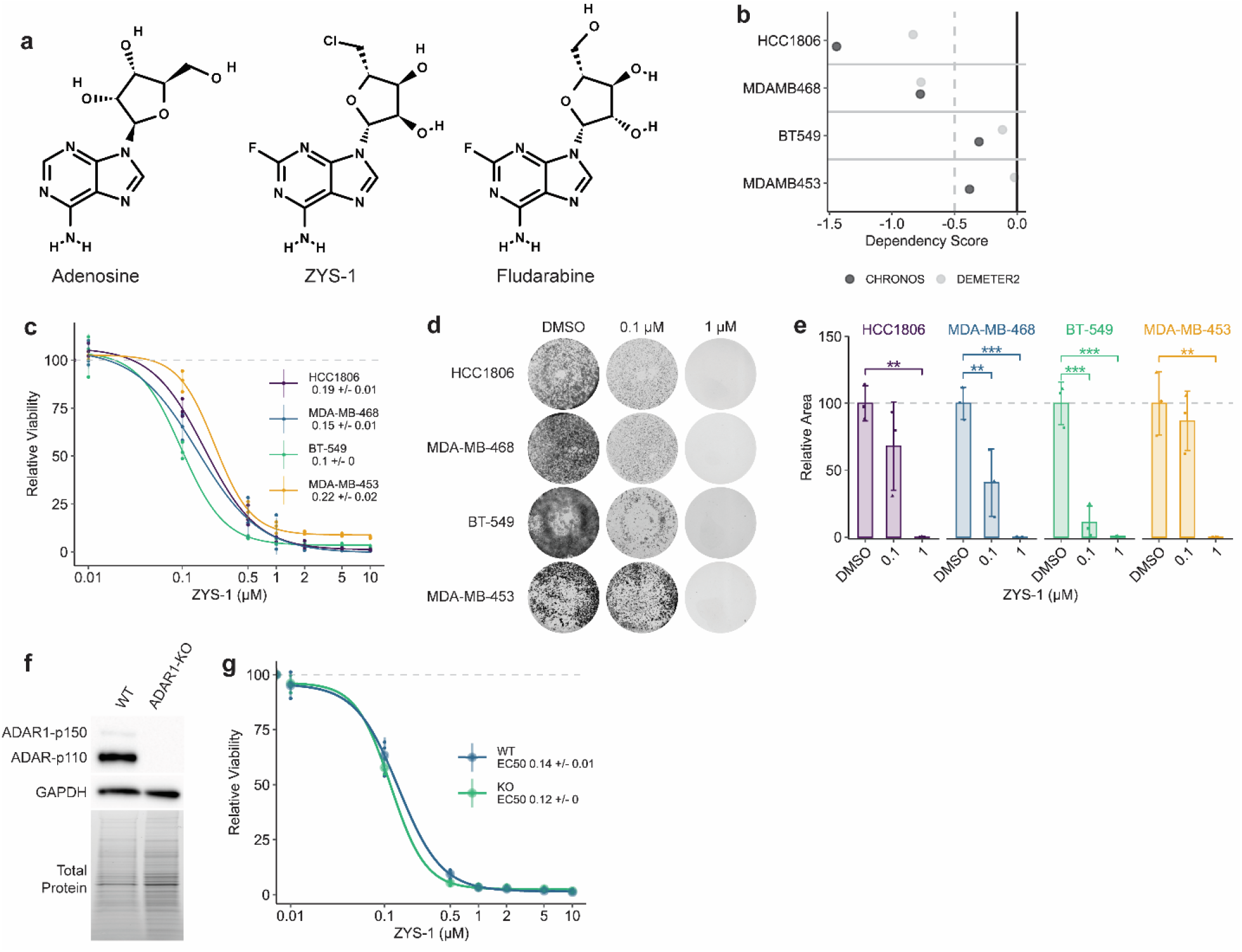
ZYS-1 is broadly cytotoxic to breast cancer cells, irrespective of ADAR1 expression or ADAR1-dependency. **a** Structures of adenosine, ZYS-1, and fludarabine, made using PubChem Sketcher V2.4 (Ihlenfeldt et al. 2009). **b** CHRONOS and DEMETER2 ADAR1 dependency scores for relevant TNBC cell lines as evaluated using CRISPR-Cas9 knockout screening or RNAi screening, respectively (Broad 2024). A CHRONOS or DEMETER2 score below –0.5 indicates gene dependency (Broad 2024). **c** Dose-response evaluation of cell viability for ZYS-1 in cell lines given in **b**, as measured using the CellTiter-Glo 2.0 assay. Representative images (**d**) and quantification (**e**) of crystal violet staining following ZYS-1 treatment of cell lines given in **b. f** Immunoblot analysis of ADAR1 protein expression in WT and ADAR1-KO BT-549 cells. **g** Dose-response evaluation of cell viability for ZYS-1 in WT and ADAR1-KO BT-549 cells, as measured using the CellTiter-Glo 2.0 assay. For **c** and **g**, large points are the average of three biological replicates, and smaller points are the average of three technical replicates performed for each biological replicate. For **e**, bars indicate the average of three biological replicates. Error bars represent the average ± SD. Statistical analysis for **e** performed using one-way ANOVA with post-hoc Tukey. ** p < 0.01, *** p < 0.001.

## Results

### ZYS-1 is broadly cytotoxic

A selective inhibitor of ADAR1 would be expected to be more cytotoxic towards ADAR1-dependent cell lines than ADAR1-independent lines. As such, we evaluated the effect of ZYS-1 treatment on cell viability in a panel of ADAR1-dependent and ADAR1-independent cell lines. We chose two dependent cell lines (HCC1806 and MDA-MB-468) and two independent cell lines (BT-549 and MDA-MB-453) based on either DepMap data (Broad 2024) (Fig. 1b) or our previous studies (Kung et al. 2021). Irrespective of the ADAR1-dependency status, ZYS-1 was strongly cytotoxic towards all cell lines evaluated (Fig. 1c). As measured by an ATP-dependent luciferase-based assay for cell viability, we found the EC50 of ZYS-1 to vary little across the cell lines in our panel and to be lowest in the ADAR1-independent cell line BT-549. Crystal violet staining of cells treated with ZYS-1 supported the robust toxicity of ZYS-1 (Figure 1d-e).

To more rigorously evaluate the specificity of ZYS-1, we assessed its effect on cell viability in WT and ADAR1 knockout (ADAR1-KO) BT-549 (Fig. 1f). The EC50 for ZYS-1 in WT and ADAR1-KO BT-549 cells was nearly identical (0.14 and 0.12 µM, respectively (Fig. 1g)). Importantly, the WT and ADAR1-KO BT-549 were equally viable in the absence of ZYS-1 (Supplemental Figure 1a). We attempted to assess the effect of ZYS-1 by crystal violet staining, but found that ADAR1-KO BT-549 cells detached easily during the staining process (Supplemental Fig. S1b). Together with the results above, these data strongly indicate that ZYS-1 does not cause cell death by affecting ADAR1.

### ZYS-1 does not phenocopy ADAR1 depletion

Depletion of ADAR1 in ADAR1-dependent cells produces a viral mimicry phenotype that includes activation of PKR (indicated by autophosphorylation on Thr446 (Romano et al. 1998)) and MDA5, with the latter driving a type I IFN response (Liu et al. 2019; Gannon et al. 2018). Both of these phenotypes were observed upon ZYS-1 treatment in the original study reporting ZYS-1 as an ADAR1 inhibitor (Wang et al. 2025). We evaluated the effect of ZYS-1 on PKR activation and ISG induction using two ADAR1-dependent cell lines that we have previously studied (Cottrell et al. 2021; Kung et al. 2021). ZYS-1 treatment of the ADAR1-dependent cell lines HCC1806 and MDA-MB-468 had no effect on PKR activation (Fig. 2a-b). Previously ZYS-1 has been shown to reduce ADAR1 expression (Wang et al. 2025), however, we observed no change in ADAR1 expression in ZYS-1-treated HCC1806 and MDA-MB-468 (Fig. 2a and Fig. 2c). Increased expression of ISGs is a phenotype of ADAR1 depletion in ADAR1-dependent cancer cell lines (Cottrell et al. 2021; Liu et al. 2019). We evaluated the expression of four ISGs in ZYS-1-treated HCC1806 and MDA-MB-468. ZYS-1 did not affect ISG expression in HCC1806 and caused elevated expression of only one ISG in MDA-MB-468 (Fig. 2d). Together with our evaluation of PKR activation, these data strongly indicate that ZYS-1 does not phenocopy ADAR1 depletion.

**Figure 2.**
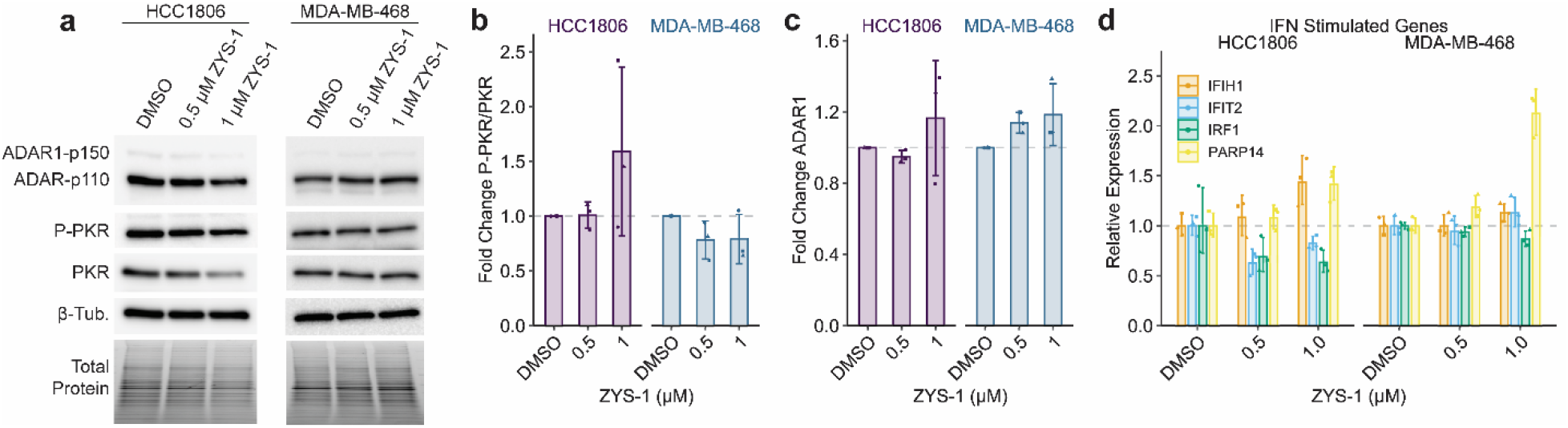
Treatment with ZYS-1 does not induce PKR activation or ISG expression. **a** Immunoblot analysis of PKR activation (P-PKR) in the ADAR1-dependent cell lines HCC1806 and MDA-MB-468 following treatment with ZYS-1. **b** and **c**, Quantification of the immunoblot shown in **a. d** qPCR analysis of IFN-stimulated gene expression in ZYS-1-treated HCC1806 and MDA-MB-468 cells. Bars indicate the average of three biological replicates. Error bars represent the average ± SD.

### ZYS-1 does not inhibit A-to-I editing

A-to-I editing by ADAR1 can be easily evaluated by sequencing of cDNA. Since inosine more readily base-pairs with cytidine, cDNA made from an edited RNA will have a guanosine at the editing site (for the sense strand of duplex cDNA or a cytidine on the anti-sense strand). Sanger sequencing is routinely used to evaluate the level of editing at a given site by comparing the peak heights of A or G (T or C on the antisense stand). We used this approach to evaluate the effect of ZYS-1 treatment on A-to-I editing in cells and in vitro using purified ADAR1 and a well-characterized substrate RNA. For editing in cells, we evaluated editing at three previously characterized editing sites. Editing at all three sites was greatly reduced by knockdown of ADAR1 in HCC1806 (Fig. 3a-b, Supplemental Figure S2). In ZYS-1-treated cells, editing was either not affected or increased for all but one site (Fig. 3c-h). For an editing site in the gene ZDHHC20, we observed significantly reduced editing at the highest concentration of ZYS-1 used in BT-549, but not in the other cell lines evaluated. It should be noted that this reduced editing occurred with 1 µM ZYS-1 treatment – a dose that is an order of magnitude higher than the EC50 for cell viability in this cell line.

**Figure 3.**
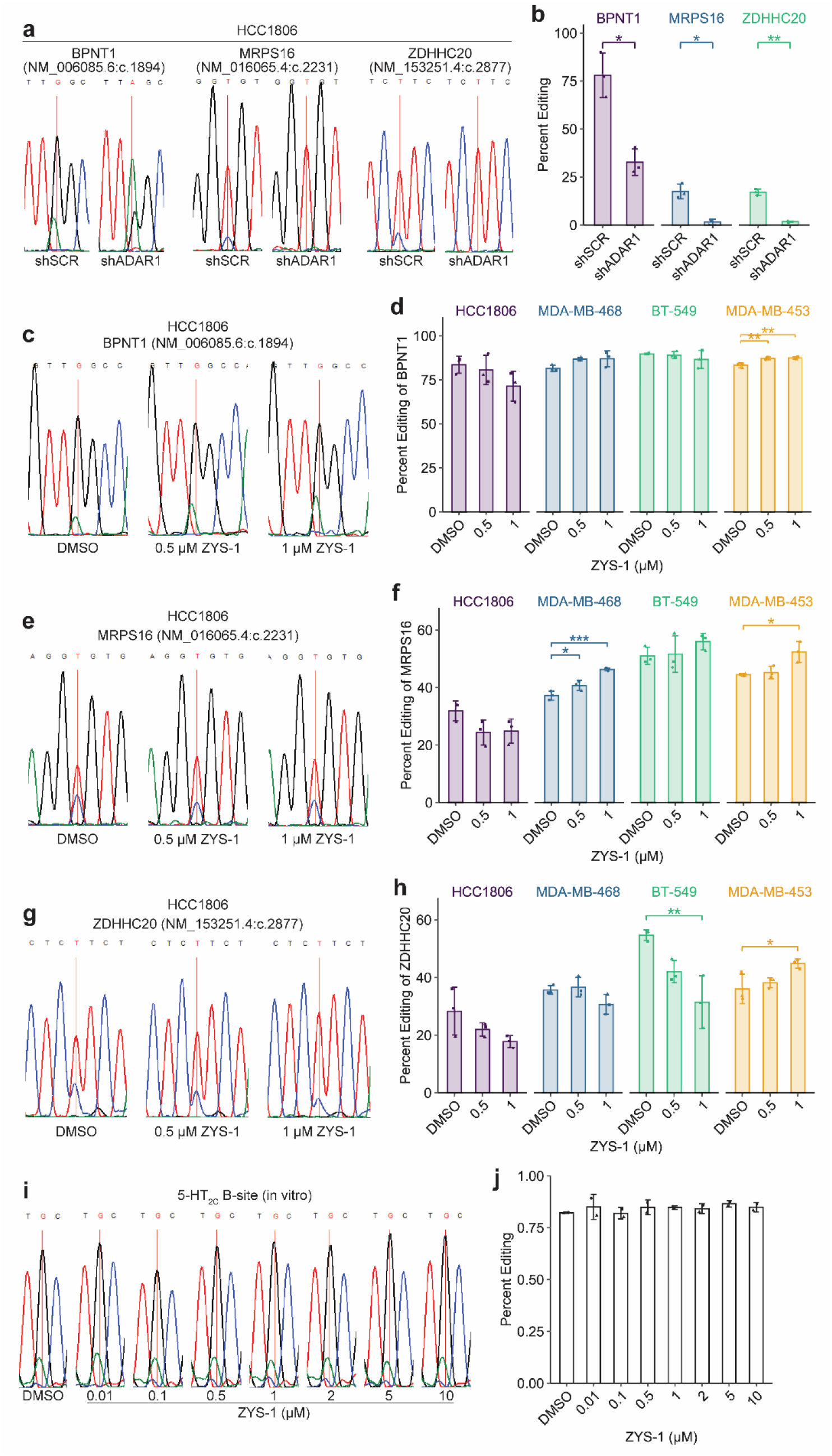
ZYS-1 treatment does not inhibit A-to-I editing by cellular or recombinant ADAR1. **a** Sanger sequencing trace of BPNT1, MRPS16, and ZDHHC20 editing sites in HCC1806 control or ADAR1 knockdown cells. **b** Quantification of percent editing using Sanger sequencing results depicted in **a**. Representative Sanger sequencing trace of BPNT1 (**c**), MRPS16 (**e**), and ZDHHC20 (**g**) editing sites in HCC1806 following treatment with ZYS-1. Percent editing quantification of the BPNT1 (**d**), MRPS16 (**f**), and ZDHHC20 (**h**) transcripts using Sanger sequencing in relevant TNBC cell lines. **i** Sanger sequencing trace of the 5-HT_2C_ B-site following incubation with purified ADAR1 and ZYS-1 as indicated. **j** Quantification of B-site percent editing using Sanger sequencing results depicted in **i**. For **a, c, e, g**, and **i**, sites that are edited by ADAR1 are indicated by the red vertical line. Bars indicate the average of three biological replicates (**b, d, f**, and **h**) or two experimental replicates (**j**). Error bars represent the average ± SD. Statistical analysis performed using Student’s t-test for **b** and one-way ANOVA with post-hoc Tukey for **d, f**, and **h**. * p < 0.05, ** p < 0.01, *** p < 0.001.

Since it is known that other proteins can regulate A-to-I editing at specific sites (Hong et al. 2018; Freund et al. 2020), it is possible that some changes in editing observed in cells upon ZYS-1 treatment could be the result of off-target effects. To evaluate direct inhibition of ADAR1, we performed an in vitro A-to-I editing assay with purified ADAR1 and an *in vitro* transcribed RNA containing a portion of the mRNA coding for 5-hydroxytryptamine receptor 2C (5-HT_2C_, encoded by *HTR2C*). The 5-HT_2C_ mRNA contains a well-characterized ADAR1 editing site, known as the ‘B-site’, and is frequently used in A-to-I editing assays (Burns et al. 1997). The 5-HT_2C_ B-site was well edited by ADAR1 in DMSO controls (Fig. 3i-j), though we observed differences in editing based on the lot of ADAR1 used (see Supplemental Fig. S3 for additional data from a different enzyme lot and additional assay controls). Addition of ZYS-1 to the editing assay had no effect on A-to-I editing at any dose used (Fig. 3i-j and Supplemental Fig. S3a-b). Together with the cellular A-to-I editing data above, these data indicate that ZYS-1 is not an ADAR1 inhibitor.

## Discussion

Many studies have now shown the therapeutic potential of targeting ADAR1 to treat a range of cancers (Liu et al. 2019; Ishizuka et al. 2019; Gannon et al. 2018; Kung et al. 2021). In addition to the cell-intrinsic effects of disrupting ADAR1, the cell-extrinsic effects, such as overcoming resistance to immune checkpoint blockade, strengthen the potential of therapies targeting ADAR1. As such, identifying small molecule inhibitors of ADAR1 is of great importance. Previously reported ADAR1 inhibitors, the adenosine analogues 8-azaadenosine and 8-chloroadenosine, have not proven to be effective in subsequent studies (Cottrell et al. 2021). Unfortunately, our data above indicate that the adenosine analogue ZYS-1 is not a selective inhibitor of ADAR1.

While we show that ZYS-1 effectively kills many breast cancer cell lines, it does so not by inhibiting ADAR1. Our data are very different from those reported in the initial publication describing ZYS-1 as an ADAR1 inhibitor (Wang et al. 2025). There are a few possible explanations for the differences between these studies. First, the original paper reporting ZYS-1 as an ADAR1 inhibitor, Wang et al., 2025, used a commercial assay for Adenosine Deaminase (ADA) to assess inhibition of A-to-I editing by ADAR1 (Wang et al. 2025). That assay is not designed to assess ADAR1 activity and has been reported to be incapable of detecting A-to-I editing (Zhang et al. 2025). Here we used a conventional in vitro A-to-I editing assay to evaluate inhibition of ADAR1 by ZYS-1 and observed no inhibition, consistent with the findings of a recent preprint (Zhang et al. 2025). Second, while Wang et al., 2025 provided evidence that ZYS-1 inhibited A-to-I editing within cells, they evaluated only one editing site in two cell lines. We evaluated three editing sites across four cell lines and observed a significant editing reduction at only one site, in one cell line. Most sites had no observable change, and we even found some sites to have increased editing. Because A-to-I editing by ADAR1 can be regulated by other proteins (Freund et al. 2020; Hong et al. 2018), it is possible that a small molecule could indirectly affect A-to-I editing, which could explain both increased or decreased editing at specific sites. Third, in Wang et al., 2025, the authors evaluated the effect of ZYS-1 on cell viability in control and ADAR1 knockout cell lines. They found ZYS-1 to be less effective in the ADAR1 knockout cells. However, in that experiment, knockout of ADAR1 greatly reduced cell viability, which could have reduced the efficacy of ZYS-1 (which we suspect is inhibiting DNA synthesis, see below). In contrast, we performed a similar experiment with control versus ADAR1 knockout cells, using an ADAR1-independent cell line to ensure that knockout did not affect cell viability. In our experiment, ZYS-1 efficacy was not affected by ADAR1 knockout.

While ZYS-1 is not a selective ADAR1 inhibitor, it is highly cytotoxic to cancer cells. ZYS-1 is a derivative of fludarabine. Fludarabine has a long history as a chemotherapeutic used for multiple leukemias and lymphomas (Gandhi and Plunkett 2002). Its mechanism of action involves inhibition of DNA synthesis, through both inhibition of DNA polymerase and ribonucleotide reductase (Gandhi and Plunkett 2002). We suspect that ZYS-1 is functioning through the same mechanism. This would explain its cytotoxicity.

Finally, it is our opinion that adenosine analogues are unlikely to inhibit ADAR1 (or ADAR2). Early studies demonstrated that ADAR1 is highly selective for dsRNA(Bass and Weintraub 1988; Wagner and Nishikura 1988; Polson et al. 1991). Unlike ADA, ADAR1 cannot be inhibited by coformycin or 8-azanebularine – both of which structurally resemble the hydrated reaction intermediate of the adenosine deamination reaction (Polson et al. 1991; Mendoza et al. 2023). However, when incorporated into a duplexed RNA, but not single-stranded RNA, 8-azanebularine potently inhibits ADAR1 with a low nanomolar IC50 (Mendoza et al. 2023). These findings suggest that additional interactions between ADAR1 and the substrate dsRNA are required to adjust the conformation of the active site and allow binding to adenosine – a model that was first suggested over three decades ago (Polson et al. 1991). This model is supported by structural studies of ADAR1 and ADAR2 indicating multiple interactions between active site adjacent residues and either the phosphodiester backbone of the dsRNA substrate or the ‘orphan’ base opposite the edited adenosine (Fisher and Beal 2024; Matthews et al. 2016; Deng et al. 2025). For ADAR2, the interactions between the deaminase domain and the dsRNA substrate cause long-distance changes to the deaminase domain structure (Matthews et al. 2016). These prior observations, along with our evaluation of ZYS-1 and other adenosine analogues (Cottrell et al. 2021), all indicate that selective inhibition of ADAR1 is unlikely to be accomplished via an adenosine analogue.

## Materials and Methods

### Cell Culture and Lentiviral Transduction

The cell lines (HEK293T (RRID: CVCL_0063), MDA-MB-453 (RRID: CVCL _0418), MDA-MB-468 (RRID: CVCL_0063), BT-549 (RRID: CVCL_1092), and HCC1806 (RRID: CVCL_1258)) were purchased from the American Type Culture Collection and cultured as described previously (Young et al. 2025). Detailed methods are provided in the Supplemental Information. For some experiments we used the above cell lines engineered for inducible expression of Cas9. These iCas9 cell lines were described previously by our laboratory (Young et al. 2025). The iCas9 lines were used for ADAR1 knockout (Fig. 1e-f, BT-549 iCas9) and for evaluating ZYS-1 efficacy in the cell lines MDA-MB-453 and MDA-MB-468. ZYS-1 (Fludarabine-Cl) was purchased from MedChemExpress (Cat# HY-145443).

Lentiviral production and transduction was performed as described previously (Young et al. 2025). Detailed methods are provided in the Supplemental Information.

### Plasmids

Oligos encoding an ADAR1-targeting sgRNA (Table S1) were cloned into lenti-sgRNA-hygro (a gift from Brett Stringer (Stringer et al. 2019), Addgene #104991; RRID:Addgene_104991) by ligating annealed and phosphorylated oligos into a digested and dephosphorylated vector. The lenti-sgC1-hygro, generated previously (Young et al. 2025), expresses a control sgRNA that targets a control genomic locus (AAVS1) and has been used previously (Zou et al. 2024; Young et al. 2025). The ADAR1 sgRNA used is from the Avana library used in DepMap (Broad 2024).

The control and ADAR1 knockdown plasmids (tet-pLKO-shSCR-puro-luc and tet-pLKO-shADAR1-puro-luc) were generated by cloning of oligonucleotides encoding either shSCR or shADAR1 into the plasmid tet-pLKO-puro-luc by ligating annealed and phosphorylated oligos into a digested and dephosphorylated vector, see Supplemental Information for oligonucleotide sequences. To generate tet-pLKO-puro-luc, first a PstI site was inserted at the end of PuroR in tet-pLKO-puro (a gift from Dmitri Wiederschain (Wiederschain et al. 2009), Addgene plasmid # 21915, RRID:Addgene_21915) by site directed mutagenesis using PrimeSTAR Max DNA Polymerase (Takara Bio) and In-Fusion Snap Assembly (Takara). A region of the plasmid pRG01-U6-DR-crRNA-BsmbI(x2)-6T; EFS-Puro-2A-Fluc-WPRE (a gift from Sidi Chen (Chow et al. 2019), Addgene plasmid # 123362, RRID:Addgene_123362) encoding PuroR-2A-Fluc was PCR amplified and digested with BsiWI and PstI prior to ligation into the tet-pLKO-puro-PstI digested with the same enzymes. See Supplemental Table 1 for all primer sequences. Both the control and ADAR1 targeting shRNAs used here have been used previously to evaluate ADAR1 function in breast cancer (Kung et al. 2025; Kung et al. 2021; Young et al. 2025; Cottrell et al. 2024b; Cottrell et al. 2021). The sequences for both shRNAs are listed in Table S1.

All plasmids were confirmed by Sanger sequencing and Nanopore whole plasmid sequencing as well as restriction enzyme digestion.

### Genetic Depletion by CRISPR-Cas9 or RNAi

To generate ADAR1 knockout in BT-549, BT-549 iCas9 cell lines were transduced with lentivirus made with lenti-sgADAR1-hygro or lenti-sgC1-hygro and selected with hygromycin. The cells were treated with 2 µg/mL doxycycline to induce Cas9 expression for at least seven days prior to maintenance in media without doxycycline for at least three days before experimentation. To establish the BT-549 ADAR1 knockout cell line, we screened clonal lines for ADAR1 knockout by immunoblot. We used the polyclonal sgC1 line as our control.

For knockdown of ADAR1, HCC1806 cells were transduced with either tet-pLKO-sshSCR-puro-luc or tet-pLKO-shADAR1-puro-luc and selected with puromycin. Knockdown of ADAR1 was initiated by doxycycline treatment at 2 µg/mL.

### Immunoblot

Immunoblot analysis was performed as previously described, with between 25 and 40 micrograms of protein loaded per lane (Young et al. 2025). Detailed methods are described in the Supplemental Information. For ZYS-1-treated cells, cells were treated with DMSO or ZYS-1 for 48 hours prior to harvesting protein. Primary antibodies used include: ADAR1 (Santa Cruz Biotechnology Cat# sc-73408, RRID:AB_2222767), PKR (Cell Signaling Technology Cat# 3072, RRID:AB_2277600), PKR Thr-446-P (Abcam Cat# ab32036, RRID:AB_777310), GAPDH (Santa Cruz Biotechnology Cat# sc-47724, RRID:AB_627678), and beta-tubulin (Abcam Cat# ab6046, RRID:AB_2210370).

### Cell Viability and Proliferation Assessment

To assess cell viability, 2,500 (HCC1806 and BT-549) or 5,000 (MDA-MB-453 and MDA-MB-468) cells were plated in triplicate per condition in 96-well plates. The cells were treated as indicated one day after plating. CellTiter-Glo 2.0 (Promega) was used to determine cell viability per manufacturer’s protocol three days after treatment.

To assess cell proliferation and viability by crystal violet staining, 25,000 (HCC1806 and BT-549) or 50,000 (MDA-MB-453 and MDA-MB-468) cells were plated per well of a 6-well dish. The cells were treated as indicated one day following plating. Cells were washed with 1x PBS and fixed in 100% methanol after seven to fourteen days of growth, depending on the cell line. The cells were then stained with 0.005% Crystal Violet solution containing 25% methanol (Sigma-Aldrich), prior to washing with water and imaging using a ChemiDoc Imaging System (Bio-Rad). Percent stained area was measured using Fiji (ImageJ).

### Cellular A-to-I Editing Analysis

Cells were treated with DMSO or ZYS-1 for 48 hours prior to RNA harvesting and purification using the Nucleospin RNA kit (Macherey-Nagel). cDNA was made from isolated RNA using LunaScript Supermix (NEB). Sections around the A-to-I editing sites in ZDHHC20, BPNT1, and MRPS16 were amplified using PrimeSTAR Max DNA Polymerase (Takara Bio) and the primers listed in Table S1. PCR products were then resolved by agarose gel electrophoresis and purified by the Monarch Gel Extraction kit (New England Biolabs). Purified products were Sanger sequenced by Azenta with the primers listed in Table S1. The QSVAnalyzer program was used to measure the peak heights of each A-to-I editing site. Percent editing was found by dividing the edited base (G or C) peak height by the sum of the edited and unedited (A or T) peak heights and multiplying by 100. The primer used to sequence the regions around the ZDHHC20 and MRPS16 editing sites yielded the antisense sequence of this region; percent editing for these sites was determined using C (edited) and T (unedited) peak heights.

### In Vitro A-to-I Editing Assay

A plasmid encoding a portion of the 5-HT_2C_ mRNA was synthesized by Twist Bioscience. This pTwist-5HT plasmid was linearized by KpnI restriction enzyme digest and purified by agarose gel electrophoresis as described above. The 5-HT_2C_ RNA was in vitro transcribed using HiScribe T7 Quick High Yield RNA (New England Biolabs). The in vitro transcribed RNA was purified using the Monarch Spin RNA Cleanup kit (New England Biolabs) and the quality of the RNA was assessed by denaturing polyacrylamide gel electrophoresis. The RNA was annealed in template annealing buffer (10 mM Tris-HCl pH 7.5, 1 mM EDTA, and 100 mM NaCl) by heating to 95 °C and cooling slowly to room temperature.

The in vitro A-to-I editing assay was modified from those used by the laboratory of Peter Beal, University of California Davis (Mendoza et al. 2023). In vitro editing assays were carried out in 10 µL reactions consisting of 1X ADAR1 reaction buffer (15 mM Tris pH 7.4, 26 mM KCl, 1.5 mM EDTA, 0.003% NP-40, 4% glycerol and 40 mM potassium glutamate), 0.5 mM DTT, 1 µg/mL yeast tRNA (Invitrogen), 0.16 U/µL RNase Inhibitor Murine (New England Biolabs), 4 nM 5-HT_2C_ RNA, 5 µM RT_primer (Table S1), 1 mM dNTPs and water to 8 µL. To this reaction mix, DMSO or ZYS-1 dissolved in DMSO was added prior to initiating the reaction by addition of 200 nM ADAR1-p150 (BPS Bioscience). The reaction was incubated at 30 °C for 15 minutes for ADAR1-p150 lot #231010-1 or 1 hr for lot #250218. The reaction was stopped by heating to 95 °C for 5 minutes. Reverse transcription followed immediately by the addition of ProtoScript II Buffer to 1X final (New England Biolabs), DTT to 10.25 mM final, 200 U ProtoScript II Reverse Transcriptase (New England Biolabs), 8 U RNase Inhibitor Murine and water to 20 µL total. The samples were incubated at 25 °C for 5 minutes, 42 °C for 1 hour and 65 °C for 5 minutes. The cDNA was cleaned using the Monarch PCR and DNA Cleanup kit. PrimeSTAR Max DNA Polymerase was used to PCR amplify the cDNA using primers EA_F and EA_R, described in Table S1. The PCR products were purified by agarose gel electrophoresis and Sanger sequenced using the EA_F primer. A-to-I editing was quantified as described above.

### Quantitative PCR

Total RNA isolation and cDNA synthesis was performed as above. Quantitative PCR (qPCR) was performed with Luna Universal qPCR MasterMix (NEB) on a QuantStudio3 system (Thermo Scientific). The primers used are listed in Table S1. For each primer, the amplification efficiency was verified to be within 90-110% allowing ‘Fold Change’ determination via the ΔΔCt. The geometric mean Ct of EEF1A1 and HSPA5 was used for normalization.

## Data Availability Statement

All analysis scripts are available at (https://github.com/cottrellka/Smoak_et_al_2025). Dependency data (DepMap_Public_24Q4+Score, Chronos, and Achilles+DRIVE+Marcotte, DEMETER2) were acquired from the DepMap portal (https://depmap.org/portal/download/custom/, RRID:SCR_017655).

## Acknowledgements

This work was supported by R00MD016946 (K.A. Cottrell), the Purdue Institute for Cancer Research (start-up funding) NIH grant P30CA023168, and start-up funding from Purdue University Department of Biochemistry and College of Agriculture.

## Author Contributions

K.A.C. and C.N.S. conceived the project. K.A.C. and C.N.S. designed the experiments. C.N.S., E.N.G., R.N.C., and K.A.C. performed the experiments. C.N.S. and K.A.C. performed the data analysis. K.A.C. and C.N.S. wrote the manuscript. All authors edited the manuscript.

