## Supplemental Information for "ZYS-1 is not an ADAR1 inhibitor"

**Contents:**

Supplemental Methods

5-$HT_{2C}$ sequence

Supplemental Table 1

Supplemental Figures S1-S3

Uncropped Immunoblots

#### Supplemental Methods

##### Cell Culture

Cell lines (HEK293T (RRID: CVCL_0063), MDA-MB-453 (RRID: CVCL _0418), MDA-MB-468 (RRID: CVCL_0063), BT-549 (RRID: CVCL_1092), and HCC1806 (RRID: CVCL_1258)) were purchased from the American Type Culture Collection. STR profiling was used to authenticate the cell lines, which were obtained between 2023 and 2024. Mycoplasma contamination testing was performed using a PCR-based method. Cells used in experiments were under twenty-five passages. The cell line HEK293T was cultured in Dulbecco's modified Eagle's medium (DMEM) (Hyclone) with 10% fetal bovine serum (BioTechne), 1 mM sodium pyruvate (Hyclone), 2 mM glutamine (Hyclone), and 0.1 mM nonessential amino acids (Hyclone). HCC1806 cells were cultured in Roswell Park Memorial Institute 1640 media (RPMI) (Corning Cat# 10-041-CV) with 10% fetal bovine serum (BioTechne). The cell line BT-549 was cultured in RPMI as above, supplemented with recombinant insulin (Gibco) at 0.78 µg/mL. MDA-MB-453 and MDA-MB-468 cells were maintained in Leibowitz L-15 media (HyClone Cat# SH30525) with 10% fetal bovine serum (BioTechne). MDA-MB-453 and MDA-MB-468 were cultured at 37°C with atmospheric $CO_2$. All other cell lines were cultured at 37°C with 5% $CO_2$.

##### Viral Production and Transduction

Lentivirus was produced by branched polyethylenimine (~25,000 Da, Sigma-Aldrich) or LipoFexin (Lamda Biotech) transfection of HEK293T cells with pMD2.G (a gift from Didier Trono, Addgene plasmid #12259; RRID:Addgene_12259), pSPAX2 (a gift from Didier Trono, Addgene plasmid #12260; RRID:Addgene_12260), and a transfer plasmid for the expression of the sgRNA or shRNA of interest. Lentivirus-containing media was collected 2 to 3 days after transfection and filtered through a 0.45 µm filter. Lentivirus transduction of cells was done in the presence of 10 µg/mL protamine sulfate (Sigma-Aldrich) or 8 µg/mL polybrene (Sigma-Aldrich). Cells were selected with puromycin at 2 µg/mL (Sigma-Aldrich) or 150 µg/mL hygromycin (InvivoGen) depending on the transfer plasmid.

##### Immunoblot

Cell pellets were lysed on ice in RIPA lysis buffer (1% Triton X-100 (Sigma-Aldrich), 50 mM Tris pH 7.4 (Ambion), 150 mM NaCl (Ambion), 0.1% sodium dodecyl sulfate (Promega) and 0.5% sodium deoxycholate (Sigma-Aldrich)) with 1x HALT Protease and Phosphatase Inhibitor (Pierce). To quantify protein concentration, the DC Assay kit (Bio-Rad) was used. The lysate supernatant was diluted in SDS Sample Buffer (125 mM Tris pH 6.8, 30% glycerol, 10% sodium dodecyl sulfate, and 0.012% bromophenol blue) and boiled at 95 °C for 7 minutes. Between 25 and 40 micrograms of protein was loaded per lane onto 4-12% TGX Acrylamide Stain-Free gels (Bio-Rad). The Stain-Free gel was transferred onto a PVDF membrane (Bio-Rad) by TransBlot Turbo (Bio-Rad) and then blocked in either 5% milk or 5% bovine serum albumin in tris-buffered saline with tween before overnight primary antibody incubation. Following primary antibody binding and a horseradish-peroxidase conjugated secondary antibody (Jackson ImmunoResearch) incubation, Clarity Western ECL Substrate (Bio-Rad) was used for band detection via ChemiDoc (Bio-Rad). Visualized bands were then normalized to total protein using Image Lab (Bio-Rad) and the Stain-Free gel image, and the bands of interest were quantified.

**5-$HT_{2C}$ sequences**

DNA sequence (5'-3') encoding a portion of the **5-$HT_{2C}$** mRNA used for in vitro A-to-I editing assays

TAATACGACTCACTATAGTGGGTACGAATTCCCACTTACGTACAAGCTTACCTAGATATTTGTGCCCCGTCTGGATTTCTTTAGATGTT
TTATTTTCAACAGCGTCCATCATGCACCTCTGCGCTATATCGCTGGATCGGTATGTAGCAAT**A**CGTAATCCTATTGAGCATAGCCGTTT
CAATTCGCGGACTAAGGCCATCATGAAGATTGCTATTGTTTGGGCAATTTCTATAGGTAAATAAAACTTTTTGGCCATAAGAATTGCAG
CGGCTATGCTCAATACTTTCGGATTATGTACTGTGAACAACGTACAGACGTCGACTGGTAACATTTGCGTTTGATCGGGTTCTggtacc

RNA sequence (5'-3') for a portion of the **5-$HT_{2C}$** mRNA used for in vitro A-to-I editing assays

GGGTACGAATTCCCACTTACGTACAAGCTTACCTAGATATTTGTGCCCCGTCTGGATTTCTTTAGATGTTTTATTTTCAACAGCGTCCA
TCATGCACCTCTGCGCTATATCGCTGGATCGGTATGTAGCAAT**A**CGTAATCCTATTGAGCATAGCCGTTTCAATTCGCGGACTAAGGCC
ATCATGAAGATTGCTATTGTTTGGGCAATTTCTATAGGTAAATAAAACTTTTTGGCCATAAGAATTGCAGCGGCTATGCTCAATACTTT
CGGATTATGTACTGTGAACAACGTACAGACGTCGACTGGTAACATTTGCGTTTGATCGGGTTCTG

-the underlined sequence is the T7 promoter
-the lowercase sequence is the KpnI site used to linearize the pTwist-5HT plasmid for in vitro transcription
-in both sequences above, the B-site adenosine is bold

**Supplemental Table 1**

| sgRNA Sequences | |
|---|---|
| **Name** | **sgRNA sequence (5'-3')** |
| sgC1 | GAGCCACATTAACCGGCCCT |
| sgADAR1-a | ACAATGGCCCCTCAAAAGCA |

| sgRNA Cloning Oligonucleotides | |
|---|---|
| sgADAR1-a_1 | caccgACAATGGCCCCTCAAAAGCA |
| sgADAR1-a_2 | aaacTGCTTTTGAGGGGCCATTGTc |

| shRNA Sequences | |
|---|---|
| **Name** | **shRNA sequence (5'-3')** |
| shSCR | CCTAAGGTTAAGTCGCCCTCG |
| shADAR1 | GCCCACTGTTATCTTCACTTT |

| shRNA Cloning Oligonucleotides | |
|---|---|
| **Name** | **shRNA sequence (5'-3')** |
| shSCR_1 | CCGGTCCTAAGGTTAAGTCGCCCTCGCTCGAGCGAGGGCGACTTAACCTTAGGTTTTTG |
| shSCR_2 | AATTCAAAAACCTAAGGTTAAGTCGCCCTCGCTCGAGCGAGGGCGACTTAACCTTAGGA |
| shADAR1_1 | CCGGGCCCACTGTTATCTTCACTTTCTCGAGAAAGTGAAGATAACAGTGGGCTTTTTG |
| shADAR1_2 | AATTCAAAAAGCCCACTGTTATCTTCACTTTCTCGAGAAAGTGAAGATAACAGTGGGC |

| Cloning Primers | |
|---|---|
| **Name** | **shRNA sequence (5'-3')** |
| tet-pLKO-puro_PstI_F | CCGCAAGCCCGGTGCCTGCAGACGCCCGCCCCACGAC |
| tet-pLKO-puro_PstI_R | GTCGTGGGGCGGGCGTCTGCAGGCACCGGGCTTGCGG |
| PuroR_F | CGAGTACAAGCCCACGGTG |
| Fluc_R_PstI | ACTACTGCAGTGTCGACTTAACGCGTGAAT |

| PCR Primers for human A-to-I Editing sites | |
|---|---|
| **Name** | **Sequence (5'-3')** |
| ZDHHC20_F | TGCTGTACTAGGAAATGACAGAGC |
| ZDHHC20_R | AACATTCTGTGATGCCTAATTTTG |
| BPNT1_F | TGCTGTGGGAGGCAAGTTAAC |
| BPNT1_R | GAGTCCGAGGCAGACAGATC |
| MRPS16_F | GAAATCGCACACTGAAATATCC |
| MRPS16_R | TTGACTCACAACCATTCTTAGGTC |

| Primers for in vitro A-to-I editing | |
|---|---|
| RT_primer | gctataacgaatactcgagggctctgatgTACCAGAACCCGATCAAACGC |
| EA_F | CCCACTTACGTACAAGCTTACCTAG |
| EA_R | GCTATAACGAATACTCGAGG |

| qPCR Primers | |
|---|---|
| IRF1 qF | AGCAAGGCCAAGAGGAAGTC |
| IRF1 qR | ACTGTGTAGCTGCTGTGGTC |
| EEF1A1 qF | TGTCGTCATTGGACACGTAGA |
| EEF1A1 qR | ACGCTCGAGAGTGAGAGGCT |
| IFIH1 qF | AGTGATTCAGGCAACATGGG |
| IFIH1 qR | GCTGGGCAACTTCCATTTGG |
| PARP14 qF | GATGTTGCTGTTGTTACCTTTCA |
| PARP14 qR | CCAGGTGGCAGGTTTTCAA |
| IFIT2 qF | CTGGGGAAACTATGCCTGGG |
| IFIT2 qR | GTGTCCACCCTTCCTCACAG |
| HSPA5 qF | GAAAGAAGGTTACCCATGCAGT |
| HSPA5 qR | CAGGCCATAAGCAATAGCAGC |

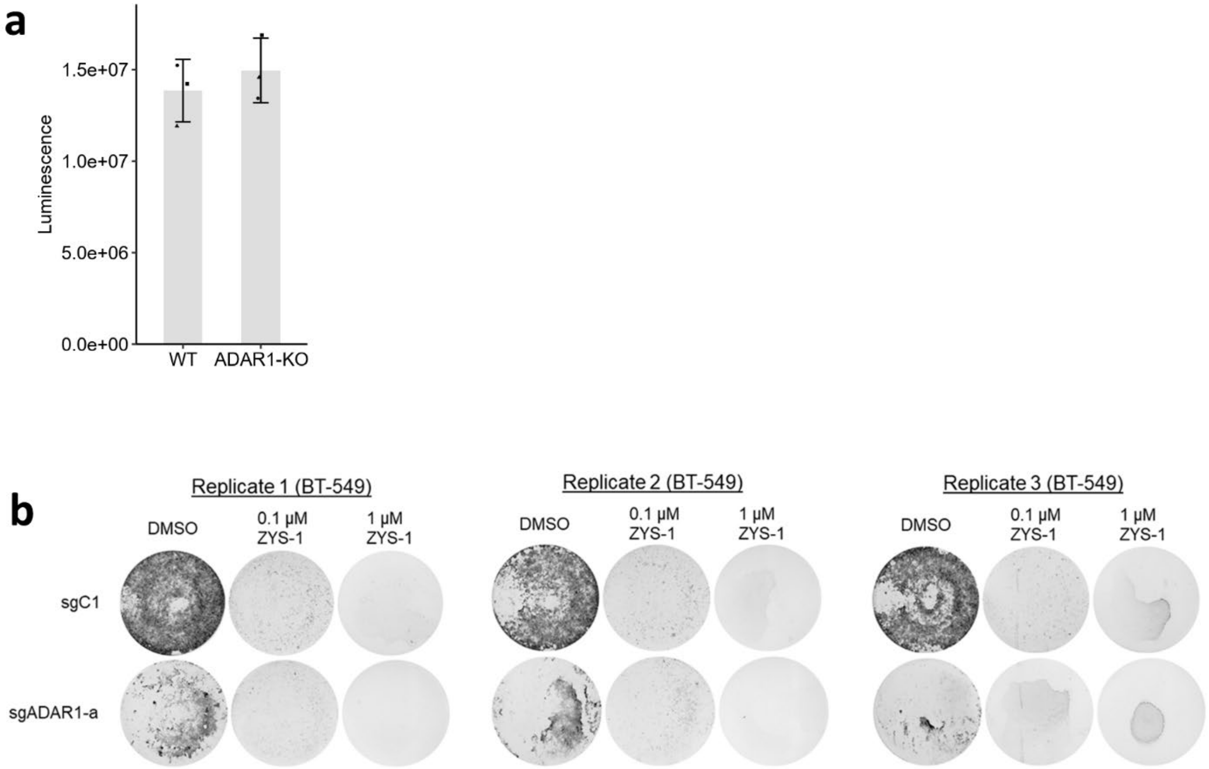


**Figure S1**

**a** Cell viability (luminescence) of DMSO treated WT or ADAR1-KO BT-549 cells as assessed by CellTiter-Glo 2.0 assay. **b** Images of crystal violet-stained foci formation assays for three replicates of WT (sgC1) or ADAR1-depleted (sgADAR1-a) BT-549 cells following treatment with ZYS-1.

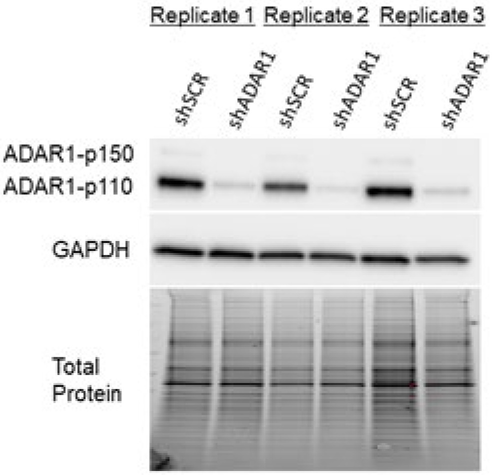


**Figure S2**

Immunoblot analysis of ADAR1 protein expression in control and ADAR1 knockdown HCC1806 cells.

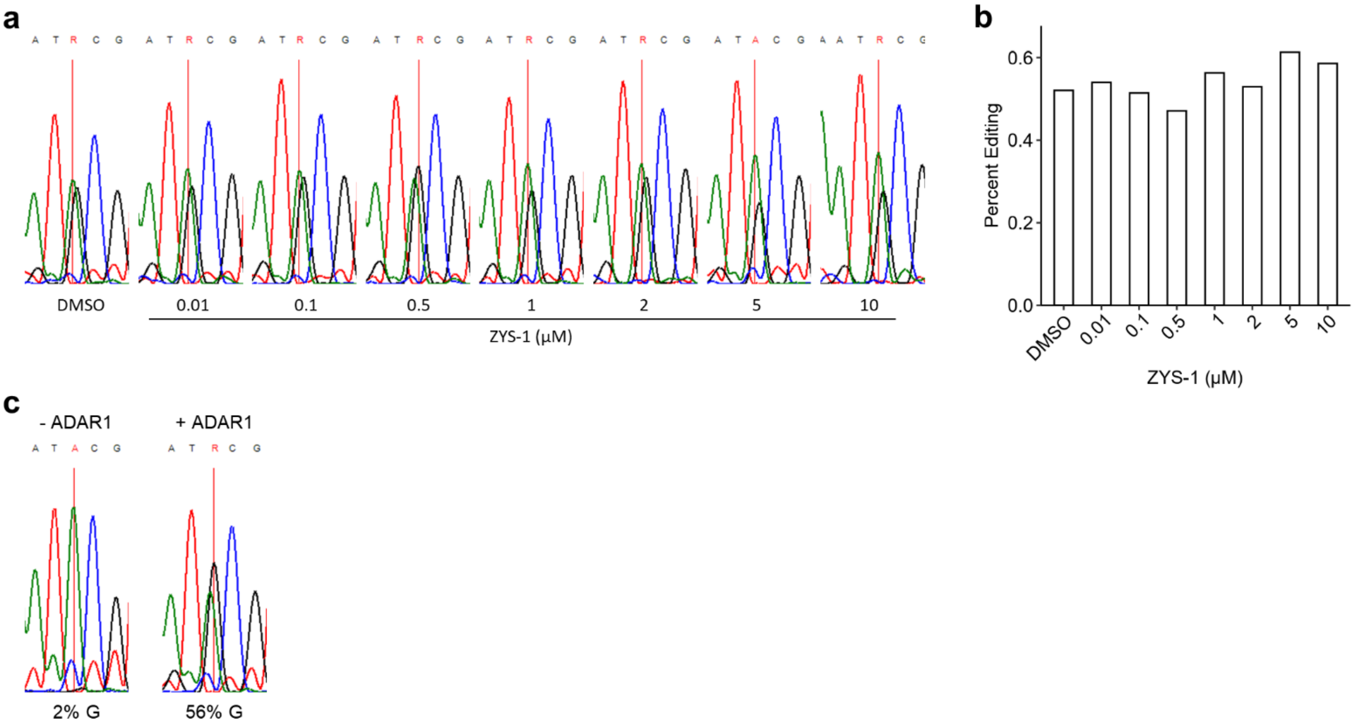


**Figure S3**

**a** Sanger sequencing trace of 5-$HT_{2C}$ B-site resulting from the in vitro A-to-I editing assay with ZYS-1 as indicated. Recombinant ADAR1 used was from a different enzyme lot than that used to generate the results in Fig. 3i-j, and incubation time was shortened from an hour to 15 minutes as well. **b** Quantification of B-site percent editing using Sanger sequencing results shown in **a**. **c** Sanger sequencing trace of the 5-$HT_{2C}$ B-site with or without the addition of ADAR1 to the in vitro A-to-I editing assay.

Below are uncropped blots for the main figures. Some blots have multiple exposures. The exposure used for each protein in the main figure is labeled in bold. Stain-free gel images for total protein are included and annotated to describe for which blots they correspond.

### FIGURE 1e.

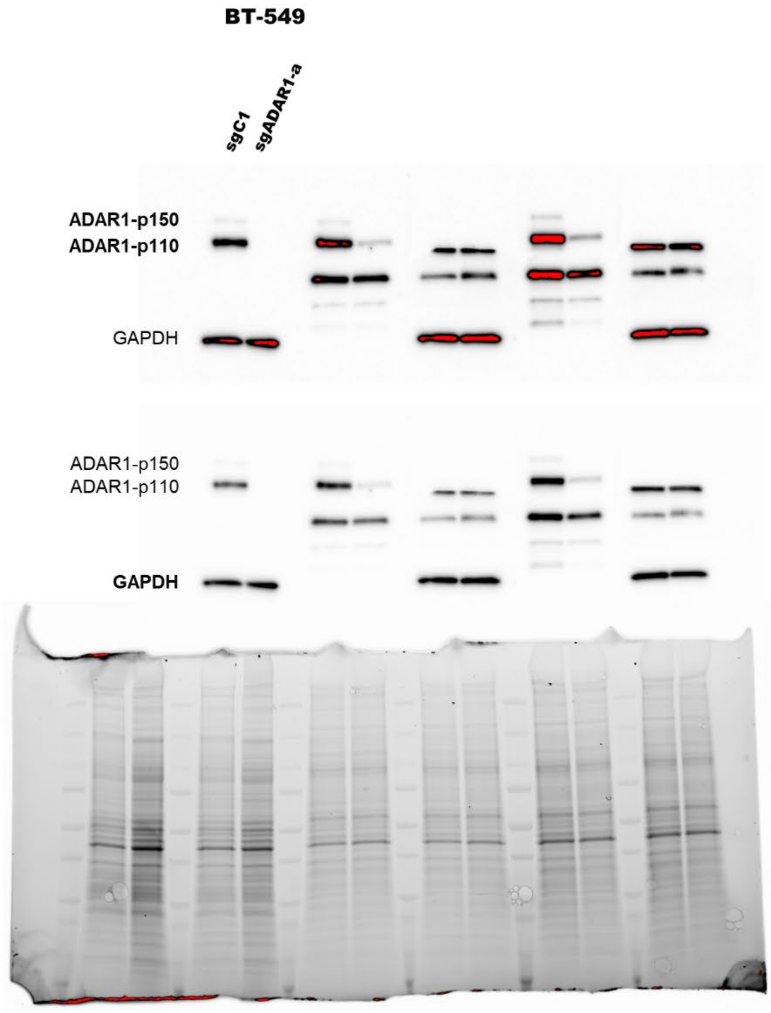


The eight rightmost samples are part of another experiment.

The two leftmost samples are part of another experiment.

### FIGURE 2a (HCC1806).

**HCC1806**

The twelve leftmost samples are replicates of this experiment that aren't visually represented.

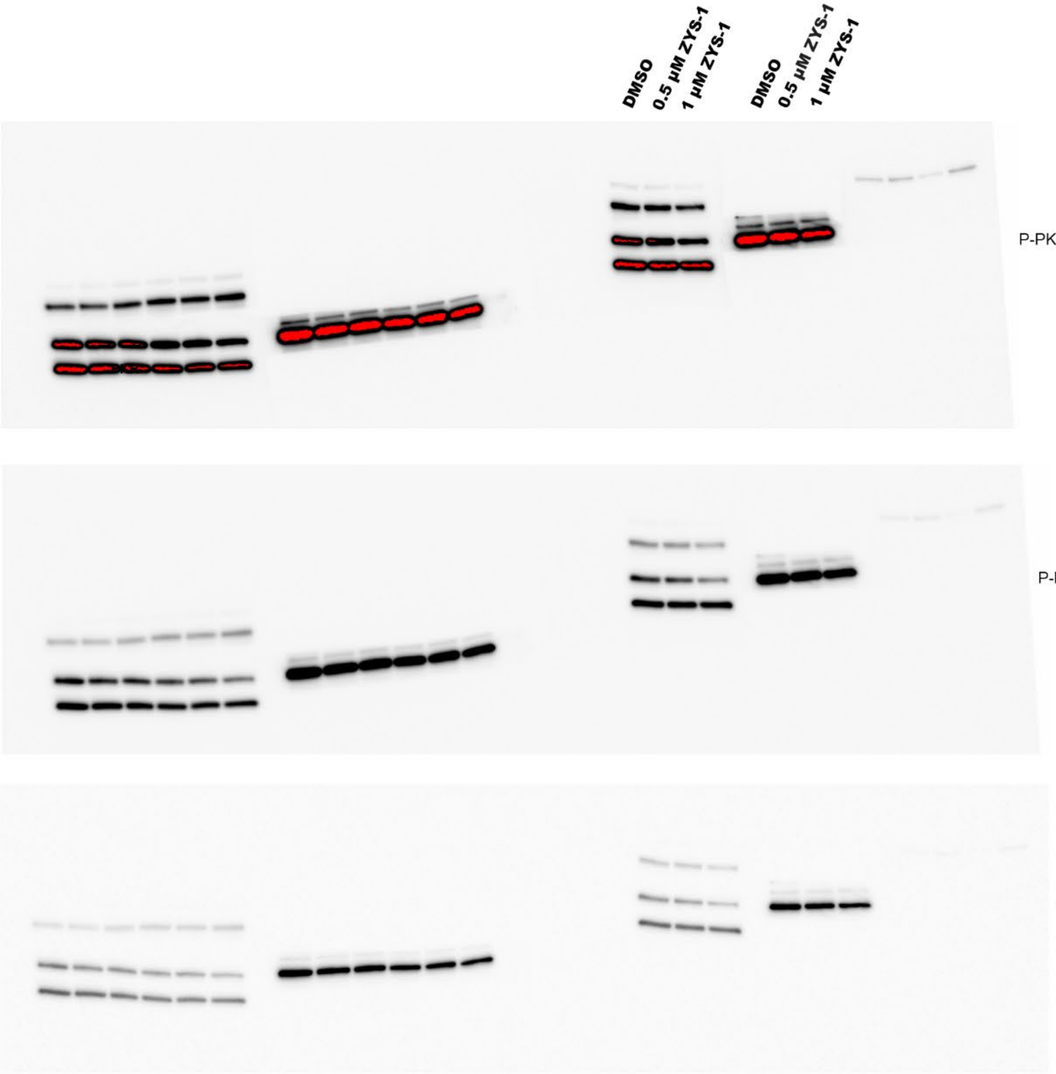


The four rightmost samples are part of another experiment.

**Total Protein**

This gel is used for the normalization of ADAR1, PKR, P-PKR, and β-tubulin.

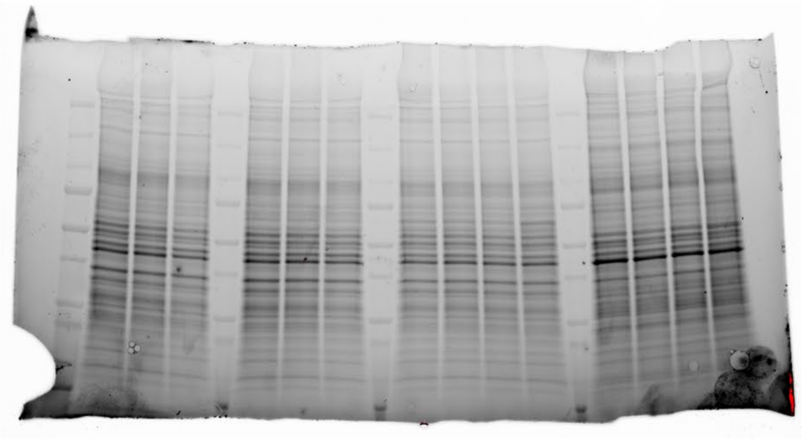

FIGURE 2a (MDA-MB-468).

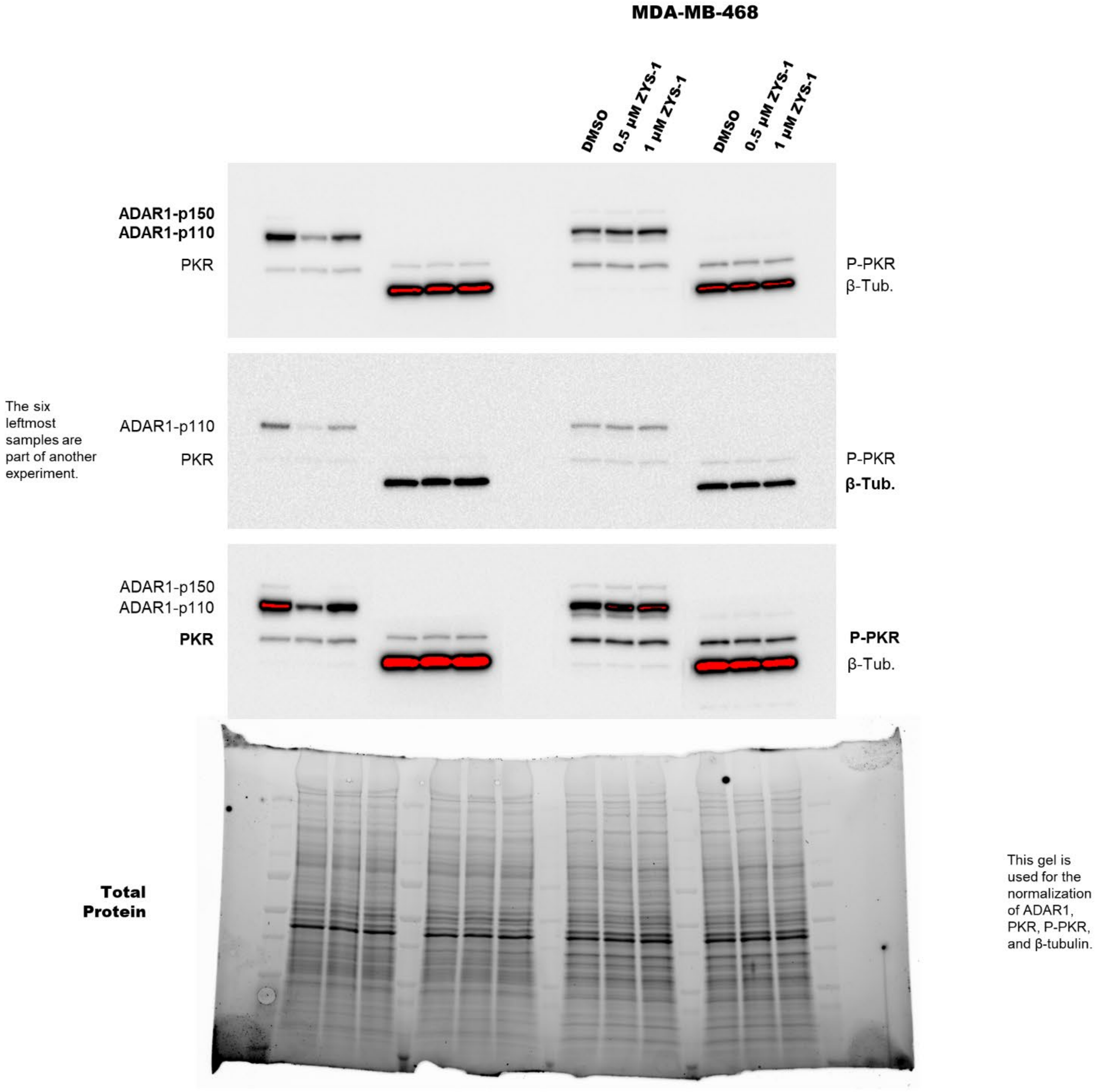

### SUPPLEMENTAL FIGURE 2

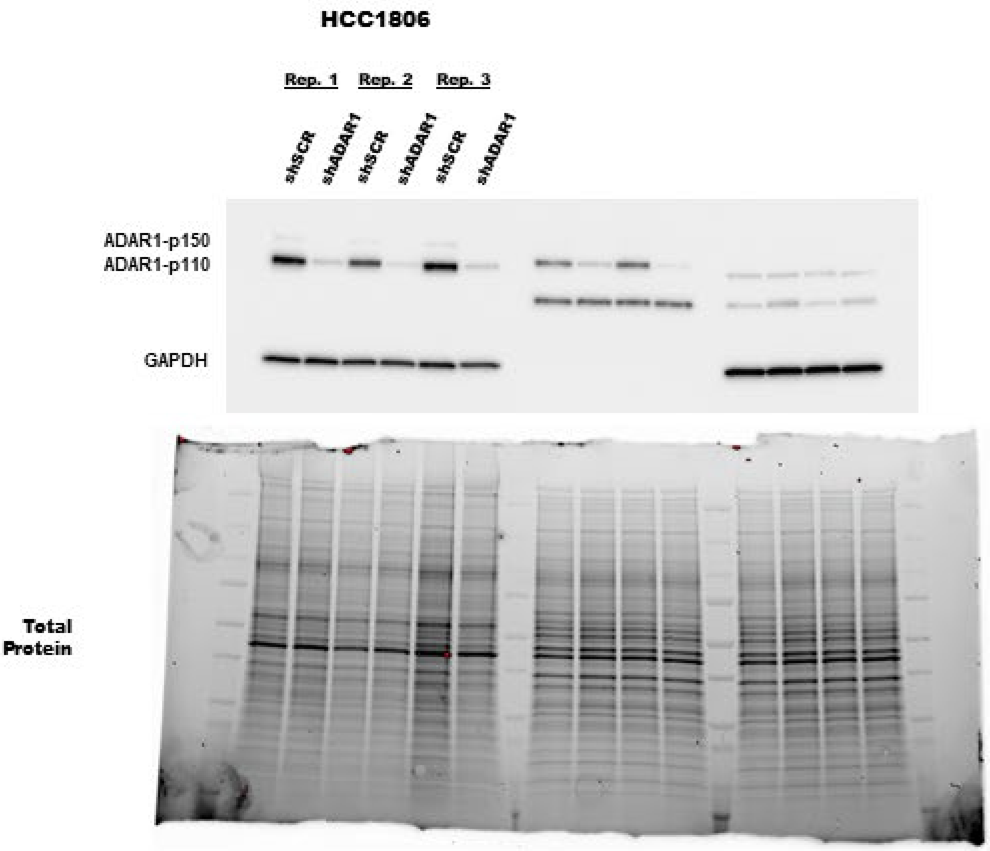


The eight rightmost samples are part of another experiment.
